# Novel candidates of pathogenic variants of the *BRCA1* and *BRCA2* genes in a 3,552 Japanese whole-genome sequence dataset (3.5KJPNv2)

**DOI:** 10.1101/2020.07.17.208454

**Authors:** Hideki Tokunaga, Keita Iida, Atsushi Hozawa, Soichi Ogishima, Yoh Watanabe, Shogo Shigeta, Muneaki Shimada, Yumi Yamaguchi-Kabata, Shu Tadaka, Fumiki Katsuoka, Shin Ito, Kazuki Kumada, Yohei Hamanaka, Nobuo Fuse, Kengo Kinoshita, Masayuki Yamamoto, Nobuo Yaegashi, Jun Yasuda

## Abstract

Identification of pathogenic germline variants yet no clinical evidence in *BRCA* genes has become important in patient care of hereditary breast and ovarian cancer syndrome (HBOC). Computational scoring and prospective cohort studies may help to identify such pathogenic variants. We annotated the variants in the *BRCA1* and *BRCA2* genes from a dataset of 3,552 whole-genome sequences obtained from members of the genome cohorts by Tohoku Medical Megabank Project (TMM) with the InterVar software. Computational impact scores (CADD_phred and Eigen_raw) and minor allele frequencies (MAF) of pathogenic (P) and likely pathogenic (LP) variants in ClinVar are used for filtration criteria. Familial predispositions in cancers among the 35,000 TMM genome cohort participants are analyzed to verify the pathogenicity. Seven potentially pathogenic variants were newly identified. Carriers of these potential pathogenic variants and definite P and LP variants among participants of the TMM prospective cohort show a statistically significant preponderance in cancer onset in sisters in the self-reported cancer history. Filtering by computational scoring and MAF is useful to identify potential pathogenic variants in *BRCA* genes for Japanese population. These results will be helpful to follow up the carriers of variants of uncertain significance in the HBOC genes.

## Introduction

Since the precision medicine initiative was launched in 2015 by the US government[1], prediction of the disease risks of individuals by using their genomic information has become plausible in a clinical setting. In Japan, gene profiling assays for cancer tissues and companion diagnostic tests for cancer-predisposing genes are now covered by the national health insurance system in Japan. These gene profiling tests can examine variations in most of the genes conferring susceptibility to two major adult-onset hereditary cancer-predisposing syndromes, hereditary breast and ovarian cancer syndrome (HBOC) and Lynch syndrome. Nowadays, the clinical significance of genetic variations of these genes is important for patient care and the health of their relatives at the bedside.

The correct judgement of germline variants in these cancer-predisposing genes is critical for the physicians who manage such patients and undertake gene profiling analyses for cancer treatment. For example, in carriers of disease-causing mutations of HBOC, prophylactic surgery is beneficial [2, 3]. Testing of *BRCA* genes may help carriers’ decision-making regarding prophylactic salpingectomy or salpingo-oophorectomy because patients with high-grade serous carcinoma arising from the fallopian tube, germline *BRCA* mutations are more prevalent in Japanese women than in other ethnic groups [4, 5]. Synthetic lethal drugs for homologous recombination defect cancers are available for patients carrying disease-causing mutations of HBOC [6, 7]. In this context, variants of uncertain significance (VUSs) would clearly be a source of major problems for clinicians. Kurian et al. reported that inexperienced breast surgeons tend to manage patients with VUSs in the *BRCA1* or *BRCA2* gene as pathogenic HBOC mutation carriers [8]. This means that lack of comprehensive annotation methods for variants might cause over-diagnosis or over-treatment in patients with *BRCA* mutations that are uncharacterized but actually benign.

To overcome these difficulties, several approaches are available. Sugano et al. reported the *BRCA1* and *BRCA2* germline variants in 135 HBOC patients and identified 28 pathogenic ones [9]. In addition, Arai et al. examined 830 Japanese HBOC pedigrees collected by the Japanese HBOC consortium and identified 49 different pathogenic variants among them [10]. Similarly, a nationwide multicenter study revealed that germline *BRCA 1/2* mutations were present in 14.7% of 634 Japanese women with ovarian cancer [5]. Lee et al. also examined the variants in the *BRCA1* and *BRCA2* genes in breast and ovarian cancer patients’ germline genomic DNA and calculated posterior probabilities for the disease-causing mutations; they identified five previously unreported variants as candidate pathogenic variants [11]. Moreover, Yost et al. reported that most VUSs in the two genes may not be pathogenic because of the low incidence of co-occurrence of allelic losses of the wild-type alleles in more than 7,000 breast and ovarian cancers with germline VUSs of the two *BRCA* genes in TCGA collection[12]. Recently, a large-scale Japanese project involving the sequencing of HBOC patients’ germline genomic DNA for 11 breast cancer predisposing genes revealed 134 pathogenic variants pathogenic germline variants concentrated in cancer patients in the *BRCA1* and *BRCA2* genes [13]. Patient-based studies for identifying germline pathogenic variants are very effective for identifying potential pathogenic variants, but cannot estimate the frequencies of those alleles in the general population. Only prospective cohorts of the general population can confirm the causality of VUSs via the collection of follow-up data and using the precise minor allele frequencies. However, in the case of follow-up surveys in prospective cohorts, it is critical to focus on the participants who need to be careful follow-up because of the limitation of the resources [14]. An appropriate method to select the participants for the detailed follow-up studies will be critical to analyze causalities of germline VUSs in the cancer predisposing genes.

Here, we describe the levels of known and potentially disease-causing variants in the *BRCA* genes among the general Japanese population, by analyzing a whole-genome reference panel for the Japanese characterized by the Tohoku Medical Megabank (TMM) Project. The TMM Project involves a combination of prospective cohort, biobanking, and genome-omics analysis (for reviews, see [15–18]). The dataset collected so far includes more than 3,500 independent whole-human-genome sequences (3.5KJPNv2) [19] with self-reported individual and family history data. We also refer to these data to test whether computational annotation can identify any variants that might cause HBOC with high penetrance.

## Materials and Methods

### Dataset

Subjects were obtained from the TMM Community-Based Cohort (TMM CommCohort) Study established by the Tohoku Medical Megabank Project [20], in which more than 120,000 adults are participating. The whole-genome sequences of some of the participants have been obtained; the criteria for selecting WGS samples are described elsewhere [21].

### Annotation of genomic variants in the *BRCA* genes

The 3.5KJPNv2 variant data were downloaded from the jMorp database (https://jmorp.megabank.tohoku.ac.jp/) [22]. The dataset is divided into two in terms of autosomal variants, namely, single-nucleotide variations (tommo-3.5KJPNv2v2-20181105open-af_snvall-autosome.vcf.gz) and indels (tommo-3.5KJPNv2v2-20181105open-af_indelall-autosome.vcf.gz), with index files. We defined the *BRCA1* and *BRCA2* regions based on GeneCards (https://www.genecards.org/) [23] as chr17:41,196,312–41,277,500 and chr13:32,889,611–32,973,809 (hg19), respectively. Variant extraction was performed with bcftools [24, 25]. The 3.5KJPNv2 VCF file integrates the multiple alleles in single lines, so the normalization was performed with bcftools.

The *BRCA* variants in 3.5KJPNv2 were annotated with the InterVar [26] command line package (default options), which depends on ANNOVAR [27]. InterVar is an analytical package to estimate the clinical impact of gene variants based on the guideline for variant interpretation by American College of Medical Genomic Guidelines and Association for Molecular Pathology in clinical sequencing [28]. InterVar includes annotations of ClinVar [29] and predicted pathogenicity, such as Combined Annotation Dependent Depletion (CADD) [30], DANN [31], and Eigen [32]. The candidate pathological variants found in the Korean population [11] were described in terms of cDNA positions. The TransVar annotation program [33] was used to obtain the genomic positions of the variants, followed by the InterVar annotation described above. To compare the variant frequencies in 3.5KJPNv2 and in gnomAD database for the *BRCA1/2* variants, we downloaded Gnomad data [34] from the associated webpage (https://gnomad.broadinstitute.org/; downloaded on 23^rd^ February, 2020). The selected variants were visualized with mutation mapper at cbioportal [35, 36].

The RIKEN 2000 genome allele frequency data [37] were downloaded from the Japanese Encyclopedia of Genetic Associations (http://jenger.riken.jp/data) and TCGA germline variant data were as described previously[38].

### Obtaining individual and family histories

TMM prospective cohort Project data are stored in a ToMMo supercomputer system, with secure data access [39]. The individual and family histories are extracted from a large data matrix consisting of self-reported findings from a paper questionnaire given to the cohort participants. The dataset consists of more than 35,000 participants in the TMM CommCohort and the data were frozen for distribution to the Japanese scientific community in 2017 as a provisional version. For most of the participants, whole-genome sequencing data are not available. The detailed method by which the participants’ histories were obtained are described elsewhere [20]. We did not use TMM Project Birth and Three-Generation Cohort data because the participants are expected to be relatively young and their family members may not be old enough to obtain positive cases [40].

The self-reported questionnaire data were filtered out for the participants who checked more than 50 items as their individual or family history that consists of around 300 entries. Most of the participants checked more than 50 items showed self-contradictory histories, such as self-history of ovarian cancer by male participants. We also checked the status regarding smoking and alcohol consumption in the potential HBOC carriers and others in the TMM CommCohort participants; we did not observe any significant difference between them (data not shown). In the statistical analysis comparing carriers of candidate *BRCA* pathogenic variants and other TMM CommCohort participants regarding self-reported individual and family histories, we employed the binomial distribution to calculate the p-value.

## Results and discussions

### Summary of *BRCA* variants in 3.5KJPNv2

More than 3,600 variants were found in the *BRCA* genes, 6.15% of which are in coding regions. The proportion of accumulated coding exonic regions of the two genes is 9.58% in hg19 and 23.1% of the total variants in the two genes in 3.5KJPNv2 are indels. Indel calling using the short-read sequence data is less reliable than single-nucleotide variants, so the indels found in 3.5KJPNv2 may require further verification using long-read sequencing data.

How many known pathogenic mutations of the BRCA genes are identified in 3.5KJPNv2? We have estimated this in a previous study on 2KJPN [41], relative to which there should be more pathogenic variants here. Table 1 indicate the annotation results of the variants in the *BRCA1* and *BRCA2* regions using the InterVar package. Ten variants in the *BRCA* genes are annotated as “pathogenic” (P) or “likely pathogenic” (LP) by referring to the ClinVar database. The accumulated frequency of pathogenic variations of *BRCA* genes in 3.5KJPNv2 is 0.0018, which might be reasonably small compared with the clinical estimation of HBOC carriers in Japan (methods).

**Table 1.** Functional annotations of the BRCA gene variants in 3.5KJPNv2.

| Data set | *BRCA1* |  | *BRCA2* |  |
| --- | --- | --- | --- | --- |
|  | 3.5KJPNv2 | GnomAD | 3.5KJPNv2 | GnomAD |
| population | 3,552 | 125,748 | 3,552 | 125,748 |
| Method | WGS | WGS & Exome* | WGS | WGS & Exome* |
| Total | 1,891 | 1,356 | 1,720 | 3,745 |
| Exonic | 83 | 879 | 139 | 2,902 |
| Clinvar P or LP (SNV) | 4 (3) | 86 (37) | 6 (2) | 228 (78) |
| InterVar P or LP (SNV) | 9 (7) | 87 (46) | 23 (14) | 275 (123) |

\* WGS: 905 individuals

To obtain more detailed insight into the 3.5KJPNv2 *BRCA* variants, we compared the results with the GnomAD database, which contains more than 130,000, multi-ethnic, human exome variants (https://gnomad.broadinstitute.org/) (Table 1). The total numbers of pathogenic variants in the *BRCA* genes are much smaller in 3.5KJPNv2 than GnomAD (Table 1). Considering the numbers of collected samples being quite different, the P or LP variants in 3.5KJPNv2 would be very similar numbers per population to that for GnomAD. The ratio of ClinVar P or LP variants are 2.81 × 10^−3^/person and 2.50 × 10^−3^/person in 3.5KJPNv2 and GnomAD, respectively. Intriguingly, there are 0 and 8 overlaps between two datasets for ClinVar and InterVar P or LP variants, respectively (data not shown). These results support the notion that pathogenic variants of a gene are highly specific to each ethnic group and therefore population specific collection of whole genome sequencing data is critical for nationwide public health care planning [18].

### Pathogenic variations in the two *BRCA* genes in the Japanese population

In the case of ClinVar, the data are based on previous reports of the identification of pathogenic variants in disease-predisposed families, so there might be new, unreported pathogenic variants to be found in the general population. To address this issue, we applied an annotation approach with InterVar. As stated above, InterVar is designed to estimate the clinical importance of human genetic variants that have not been reported previously, in accordance with the ACMG guidelines of secondary findings in clinical sequencing [28]. Interestingly, the package annotates another 13 variants as P or LP in the *BRCA* genes, as well as all of the 10 ClinVar P and/or LP variants. Among the 13 newly annotated P or LP variants, 4 are frameshift indels and 9 are nonsynonymous variants. None of these four frameshift indels is annotated with dbSNP, so it should not be considered as discordant with ClinVar. Four nonsynonymous variants detected by InterVar are annotated as “conflicting interpretation of pathogenicity” in the ClinVar database. One of the LP variants with InterVar, *BRCA1* p.L52F, shows quite high minor allele frequency (MAF) in 3.5KJPNv2 (0.0037) compared with other definite ClinVar P or LP variants. This variant is estimated to be a VUS in the Japanese HBOC consortium study [10] and “likely benign” by Lee et al. in a Korean prospective study on breast cancer patients.

There is a large publicly available dataset of Japanese whole-genome sequencing data from RIKEN [37]. It consists of deep sequencing data from 2,234 whole genomes (average depth of 25×), 1,939 of which are from BioBank Japan (BBJ), a large biobank of patients suffering from more than 50 diseases [42]. The detailed composition of the samples from BBJ is not available, but 1,276 patients with six diseases including breast cancer are included. Hence, it can be expected that pathogenic variants found in 3.5KJPNv2 might be enriched in the RIKEN dataset, although the selection criteria of the samples for the RIKEN project are unknown. As expected, two InterVar P or LP variants, *BRCA2* c.5573_5577C and *BRCA1* p.L63X, are enriched (9.75- and 16.2-fold, respectively) in the RIKEN dataset (Supplementary Table 1). In addition, a pathogenic variant not found in TMM 3.5KJPNv2 was identified (*BRCA2* p.E2877X). In contrast, the prevalent InterVar LP variant, *BRCA1* p.L52F, is not enriched in the RIKEN Japanese whole-genome dataset (0.5-fold, Supplementary Table 1). Similarly, we checked the germline variants of the *BRCA* genes in TCGA dataset [38] and found three *BRCA2* pathogenic variants that overlapped with 3.5KJPNv2 (p.T219fs, p.T1858fs, and p.N2134fs); all three of these are highly enriched in TCGA (382-fold, 318-fold, and 95.5-fold, respectively).

These results suggest that the annotation by InterVar may include false positives as well as false negatives, although no VUSs identified by the Japanese HBOC consortium are included in our estimation [10]. Precise data on the minor allele frequencies obtained by the unbiased selection of panel constituents from the general population are critical for estimating the pathogenicity of VUSs and should be included in the criteria of pathogenicity for adult-onset hereditary disorders such as HBOC based on the InterVar annotation.

### Estimate of computational scoring tools with 3.5KJPNv2 *BRCA* variants

InterVar annotates the variants’ functional impact based on the ACMG guidelines and it largely depends on previous reports to define the parameters for scoring. For example, criterion PS1 of InterVar states that “Same amino acid change as a previously established pathogenic variant regardless of nucleotide change.” This means that one needs previous knowledge about pathogenic variants to annotate a variant as “pathogenic” by InterVar. In contrast, only one supportive item, PP3, is used from computational estimations in InterVar: “Multiple lines of computational evidence support a pathogenic effect on the gene or gene product (e.g., conservation, evolutionary, splicing impact).” Hence, InterVar may underestimate the clinical impact of potentially pathogenic variants about which previous information is not available. Therefore, we would like to test whether the unbiased, computational estimations of the variants can be used to find potentially pathogenic variants without previous knowledge.

The Pearson correlation coefficients of CADD_phred with DANN_rankscore and Eigen_raw are 0.815 and 0.860, respectively, showing that both DANN and Eigen correlate well with CADD (Fig. 1a and b). However, interestingly, the distributions of ClinVar and/or InterVar P or LP variants are quite different (Fig. 1). CADD_phred and DANN_rankscore show wider distributions in P or LP variants compared with CADD_phred and Eigen_raw (Fig. 1). Interestingly, in both of the scatter plots, *BRCA1* p.L52F, a benign variant annotated as LP by InterVar, localized at a similar position to the other P or LP variants in both scatter plots in Fig. 1 A and B. Table 2 shows the details of the computational scoring in the ClinVar/InterVar P or LP variants. The ClinVar P or LP variants clearly show higher average and minimum scores for CADD_phred and Eigen_raw scores for InterVar P or LP variants, but not for DANN_rankscore. Based on this observation, we decided to use CADD_phred and Eigen_raw for further filtration of potential pathogenic mutations.

**Fig. 1.**
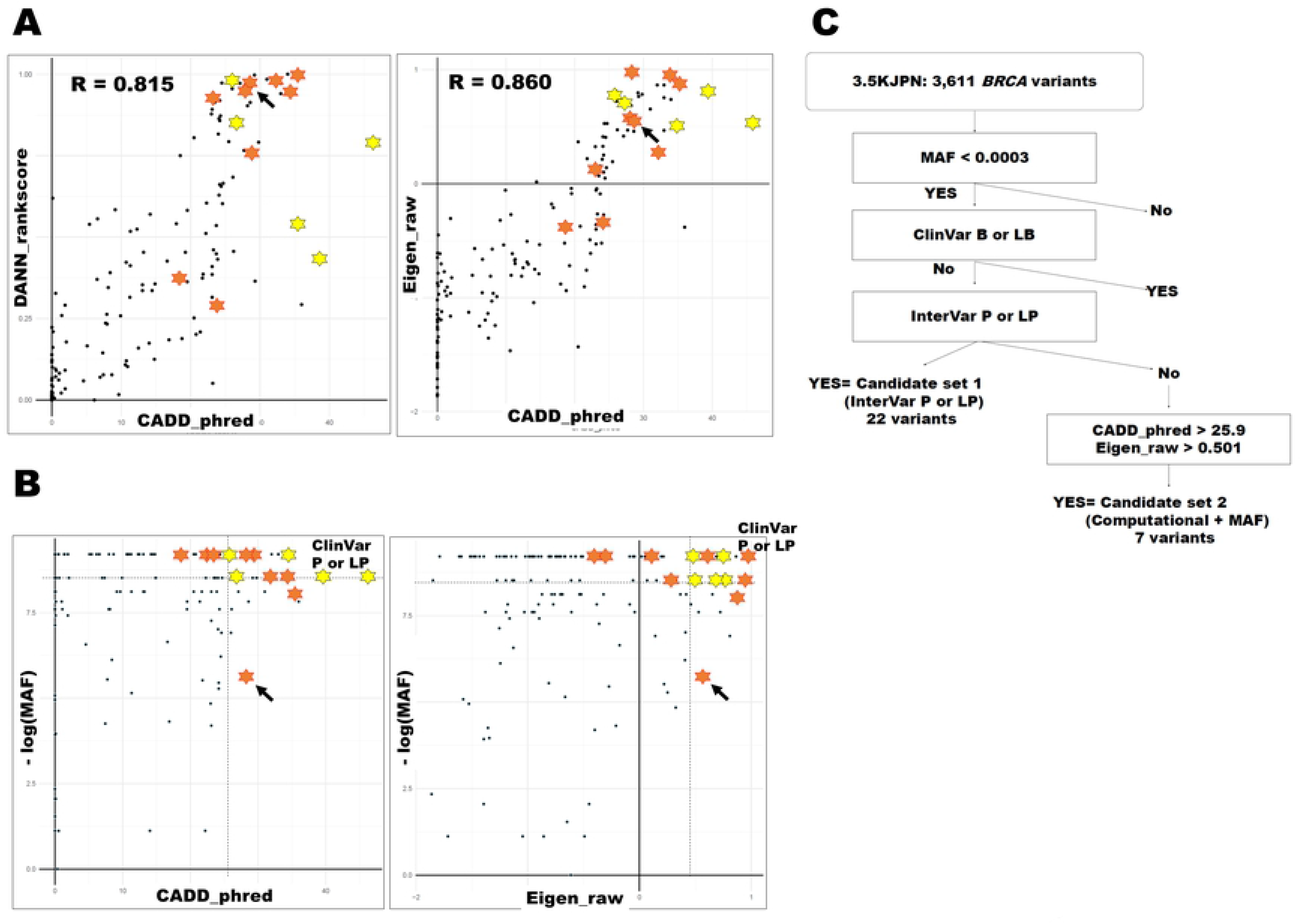
Relationships between the variant impact scores for the BRCA genes in 3.5KJPNv2. Panel a. Relationships between the CADD_phred and other two variant impact scores for the *BRCA* genes in 3.5KJPNv2. Horizontal axes indicate CADD_phred scores. Scatter plots of DANN_rankscore (left) and Eigen_raw (right) for the *BRCA* variants in 3.5KJPNv2. Yellow stars are ClinVar P or LP variants. Orange stars are InterVar P or LP variants. One InterVar variant with high MAF (0.0037) is indicated with an arrow. Panel b. Relationships between the minor allele frequencies and CADD_phred or Eigen_raw scores for the BRCA genes in 3.5KJPNv2. Vertical axes indicate inverted logarithms of minor allele frequencies for the *BRCA* variants in 3.5KJPNv2. Scatter plot of CADD_phred (left) and Eigen_raw (right) are shown. The dotted lines are maximum MAF and minimum scores of the ClinVar P or LP variants (the area defined by the threshold is indicated as “ClinVar P or LP” in the graph). Panel c. Schematic diagram of filtering steps for candidate pathogenic variants in the *BRCA* genes. Detailed filtering process is described in the main text. MAF indicates minor allele frequency.

**Table 2.** Comparison of scores for pathogenic variants in the BRCA enes in 3.5KJPN.

| InterVar P or LP | ClinVar P or LP |
| --- | --- |

|  | scored 14, not scored 9 |  |  | scored 5, not scored 5 |  |  |
| --- | --- | --- | --- | --- | --- | --- |
|  | Max | Mean | Minimum | Max | Mean | Minimum |
| CADD_phred | 46.0 | 30.3 | 18.3 | 46.0 | 34.6 | 25.9 |
| DANN_rankscore | 1.00 | 0.772 | 0.291 | 0.984 | 0.722 | 0.435 |
| Eigen_raw | 0.965 | 0.490 | -0.385 | 0.807 | 0.653 | 0.501 |
| Allele Frequency* | 3.70.E-03 | 3.22.E-04 | 1.00.E-04 | 3.00.E-04 | 1.80.E-03 | 1.00.E-04 |

\* All pathogenic or likely pathogenic variants including indels that have no computational scores

Minor allele frequencies are also critical parameters for interpreting the clinical impact of germline variants. Fig. 1B shows a scatter plot of the MAF and CADD_phred or Eigen_raw of the *BRCA* variants in 3.5KJPNv2. As expected, both CADD_phred and Eigen_raw show weak positive correlations with the reverse logarithmic minor allele frequencies (Pearson correlation coefficients = 0.172 and 0.161, respectively). The InterVar P or LP variants are localized in the vicinity of the ClinVar P or LP variants, with the exception of the *BRCA1* L52F variant (Fig. 1B). Based on these comparisons, we defined computational thresholds for possible pathogenic *BRCA* single-nucleotide variants as follows: CADD_phred ≧25.9, Eigen_raw≧0.501, and MAF≦0.0003.

Eight *BRCA* variants that fulfil these three criteria defined by the ClinVar P or LP variants are present in 3.5KJPNv2 (Supplementary Table 2). One of these, *BRCA2* p. G1529R, is annotated as “benign” or “likely benign” by ClinVar and InterVar, respectively. This variant is quite rare but found in two different ethnic groups, namely, African-Americans and non-Finnish Europeans (minor allele frequencies of 0.0003 and 0.0007, respectively; Supplementary Table 2). Because ClinVar annotated the variant as “likely benign”, we excluded it from further analysis. A summary of the filtering criteria is shown in Fig. 1C.

We also tested these criteria to the 134 potential pathogenic *BRCA* variants for women that are enriched in breast cancer cases in a previous study [13]. Among them, only 13 variants are found in the latest version of Japanese whole genome reference panel (4.7KJPN: jmorp database: https://jmorp.megabank.tohoku.ac.jp/202001/) and all of the available MAFs are ≦ 0.0003. Eighty-seven variants are P or LP for both Clinvar and InterVar and 130 variants are annotated as P or LP either ClinVar or InterVar. Three variants among 4 other are not annotated as “benign” or “likely benign” and high CADD_phred (24-35) and Eigen_raw scores (0.571-0.871) (supplementary Table 3). One exception is *BRCA1* p.K1095E, that is annotated as “likely benign” by InterVar and the both CADD_phred and Eigen_raw do not reach at our criteria. So, our criteria well corresponds to the previous studies.

A summary of the variants identified in this study is shown in Supplementary Table 2 and the distribution of the candidate pathogenic variants in the *BRCA* proteins is shown in Fig. 2. The nonsynonymous variants tend to localize at the C-terminal of the genes, while the frameshift indels and stopgains are localized between the N-terminal and the middle of the protein sequence. *BRCA2* I2675V is known as a “splicing error-causing variant” [43] and it is the most C-terminal-end variant causing large structural changes in the BRCA2 mRNA in our collection. We obtained additional annotations at the C-bioportal to draw a schematic diagram; three candidate pathogenic variants identified based on the three criteria are annotated as likely oncogenic, as well as four InterVar P or LP variants (Supplementary Table 2). This indicates that our approach can effectively identify the pathogenic variants in the *BRCA* genes. Sugano et al. described BRCA2 Y1853C as a VUS, although both ClinVar and InterVar annotated it as LP [9]. Later, Kawatsu et al. showed the pathogenic potential of this variant by experimental and genetic analyses [44]. Similarly, another variant, BRCA2 p.G2508S, is annotated as “likely neutral” by the OncoKB database. However, this variant was recently described as “moderately oncogenic” by Shimelis et al., based on a genome-wide association study of more than 12,000 cases and controls [45]. Therefore, we decided to include this variant for further study.

**Fig. 2.**
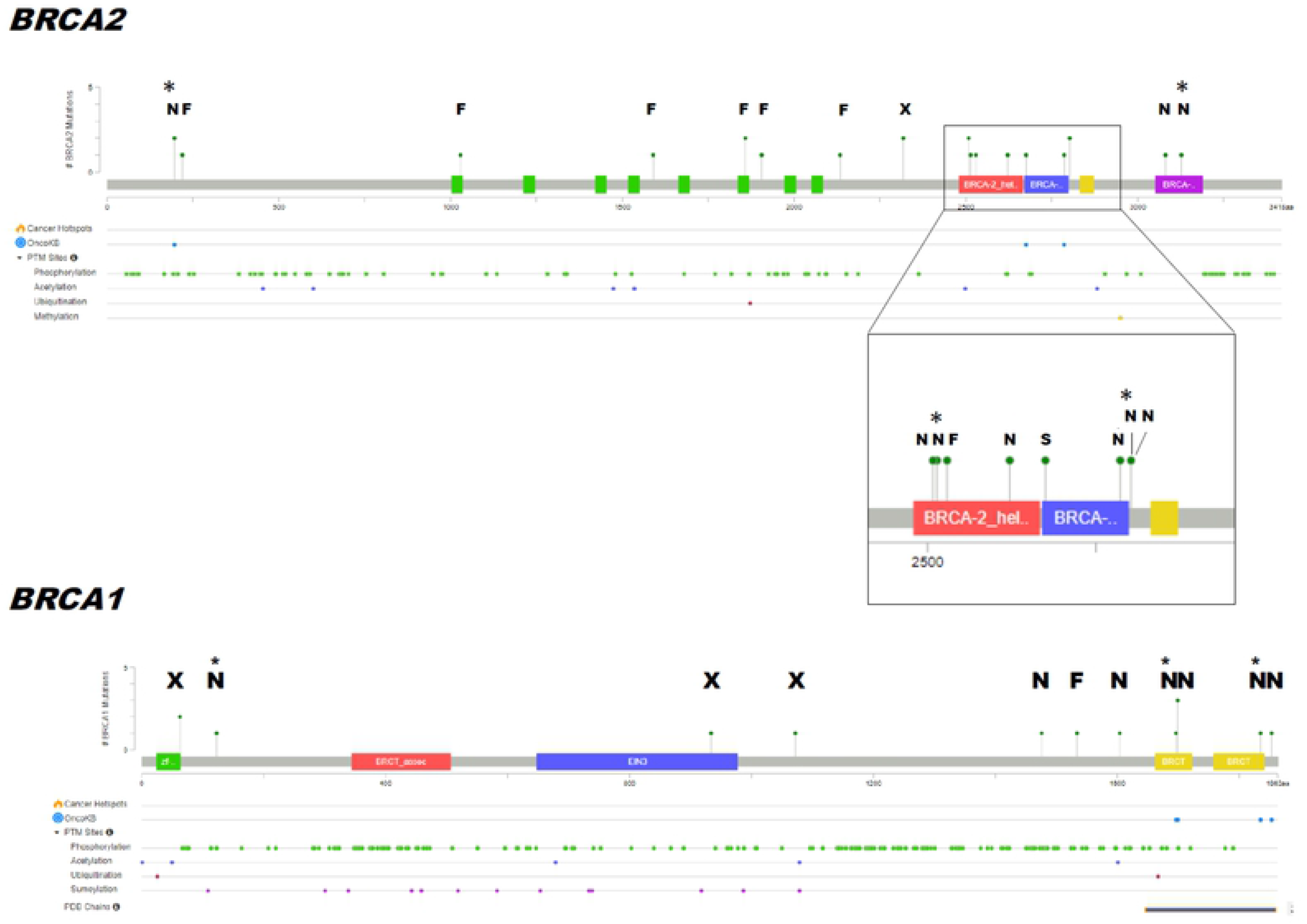
Distribution of candidate pathogenic variants of BRCA cDNA in 3.5KJPNv2. Schematic diagram of the *BRCA1* and *BRCA2* cDNA generated with Mutation Mapper on the Cbioportal. “F,” “N,” “S,” and “X” indicate frameshifts, nonsynonymous single-nucleotide variants, splicing error variants, and stopgains, respectively. The height of lollypops indicates the number of cases found in 3.5KJPNv2. Asterisks indicate variants in the computational + MAF set.

### Potentially pathogenic *BRCA* variant carriers tend to have cancer-prone family histories

Members of the TMM CommCohort reported their individual and family histories of various disorders including cancers by completing a paper-based questionnaire. It is possible that the *BRCA* pathogenic variant carriers and their family members would suffer from cancers more often than other cohort participants and their family members as most of the latter group would not carry any pathogenic *BRCA* variants. Fig. 3 indicates the numbers of cases of cancer among the participants themselves, their family members, and their spouses. Although the numbers of non-carrier participants were more than 1,500 times larger than the InterVar P or LP and computational + MAF selected *BRCA* variant carriers, the overall profiles of cancer onset were similar. For example, fathers of the participants suffered more from cancers than mothers, regardless of the participants’ status in terms of *BRCA* variants. A prominent difference between those definitely carrying potentially pathogenic *BRCA* variants and the rest of the cohort is in the ratio of cancer-bearing sisters: the InterVar P or LP carriers have a much higher rate of cancer-bearing sisters than the rest of the cohort (Fig. 3 and Fig. 3 inset; p = 3.08 × 10^−5^, chi-square test with Yates’ correction). In addition, the difference of the ratio of cancer-bearing offspring was higher in the InterVar P or LP carriers than in the others, although this was not statistically significant (Supplementary Table 4). Interestingly, the cancer onset of the participants themselves did not differ markedly between the InterVar P or LP carriers and the rest of the cohort participants. This may be reasonable as nearly half of the InterVar P or LP carriers are male and they may not suffer from *BRCA*-related cancers (Fig. 3 inset).

**Fig. 3.**
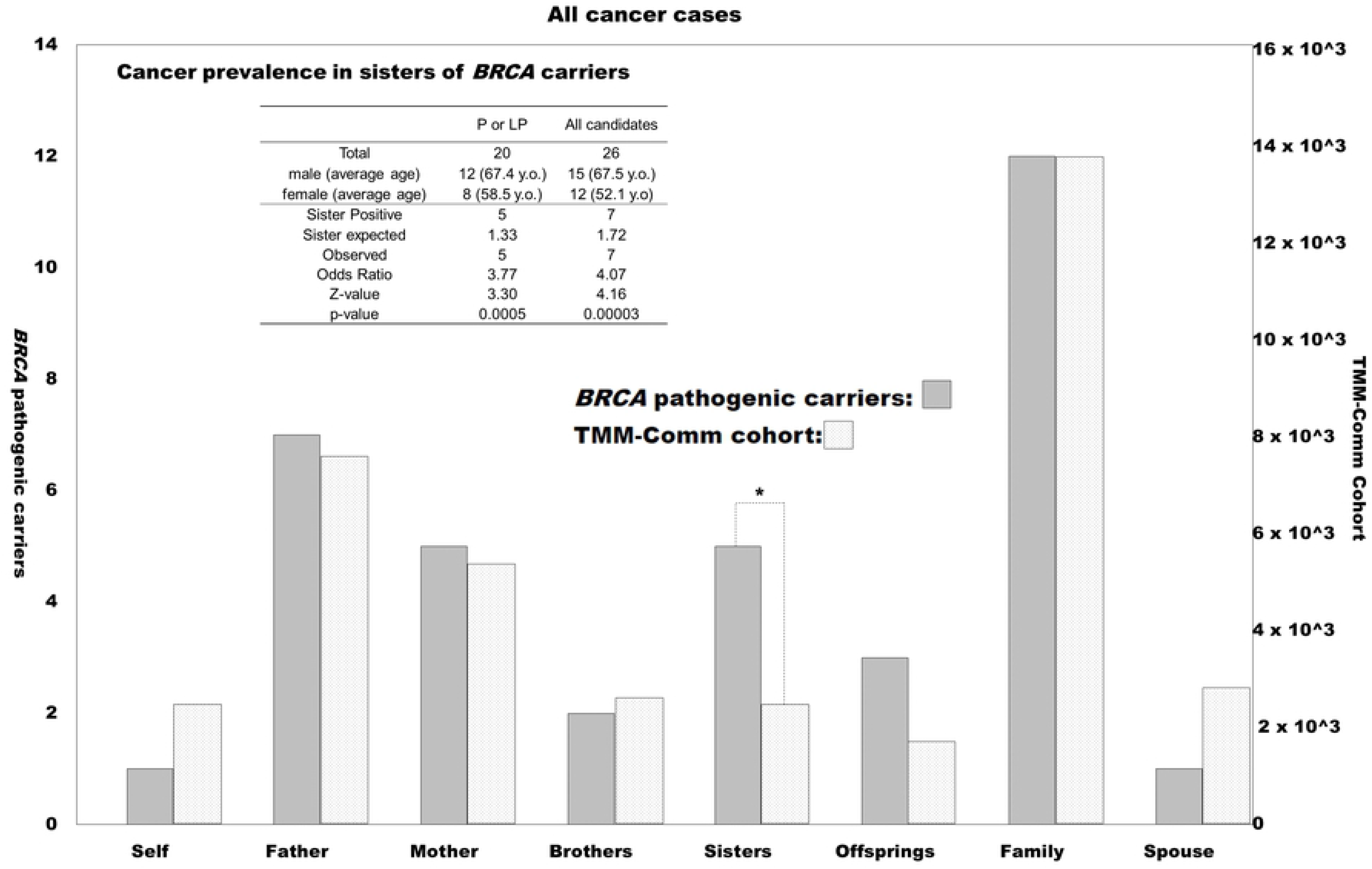
Preponderance of individual and family histories of cancer for the TMM CommCohort. Numbers of positive cases of self-reported individual and family histories of cancer in the TMM CommCohort participants. Vertical axes indicate the number of cases with positivity for each item below the horizontal axis. The right and left axes indicate the *BRCA* candidate pathogenic variant-positive and -negative cases, respectively. The scale of the vertical axes are adjusted as showing the “Family” bars same height. “Total cases” indicates the number of cases analyzed, “Self” indicates individual past history of any malignancies. “Father,” “Mother,” “Brother,” “Sister,” “Offspring,” and “Spouse” indicate the cancer-related histories of the participants’ family members. “Family” indicates any of the blood relatives for any cancers, except the participants themselves. Cases in which the “Spouse” was positive are not included in “Family.” Solid and gray bars represent numbers of cases positive for the *BRCA* candidate pathogenic variants and the rest of the TMM CommCohort cases, respectively. Asterisk indicates a statistically significant difference upon comparing with the total analyzed TMM CommCohort cases (Fig. 3 inset: Statistics of cancer prevalence in sisters of *BRCA* candidate pathogenic variants in TMM CommCohort).

Recent progress in bioinformatics may open up a completely different path for filtering the VUSs in hereditary disorders, namely, artificial intelligence-mediated approaches. One example of this is CADD, which was reported in 2014 [30]. CADD scores are calculation of all of the possible 84 billion single-nucleotide changes in the human genome. The calculation is based on machine learning using the evolutionarily conserved “proxy-neutral variants” found in both apes and humans and the recently emerged “proxy-pathogenic” rare variants in the human genome only [46]. In 2016, a further dataset, Eigen, was released, for which calculation was performed without training data but with a principal component that gives the largest diversity among the variants prepared from all possible single-nucleotide changes in the human genome [32]. These annotation tools show some clinically significant findings in genome-wide association studies (for examples, see [47, 48]). Recently, The ICGC/TCGA Pan-Cancer Analysis of Whole Genomes Consortium applied CADD for their estimation of biological impacts of cancer mutations [49]. Therefore, computational scoring is very suggestive in predicting the clinical impact of single-nucleotide variants in cancer-predisposing genes. The present study shows the potential for applying this approach to find pathogenic variations in cancer-predisposing syndromes by using genome reference panels with precise MAF estimation. This study shows that the MAF estimate with general population is much more useful for the annotation of pathogenic variants than the biased collection of population samples.

Recently, Findlay et al. reported that saturation genomics could characterize more than 4,000 *BRCA1* variants for the functionality with saturation genomics [50]. The interpretation of the variants with this data, 12 *BRCA1* variants were found in the 3.5KJPNv2. Among them, two discordant variants are found between Findlay et al. and present study. *BRCA1* p.R71S is not picked up by our survey but annotated as “loss of function” in the dataset by Findlay et al. and *BRCA1* R1699Q is picked up by our survey but annotated as “functional” in the same dataset. It can be possible that pathogenicity of *BRCA* variants would be affected by other genetic modifiers and/or environmental factors and follow-up of the carriers of these variants in prospective cohort studies may provide the clue to solve the discordance.

There was a significant preponderance of cancer in the family histories of those with potentially pathogenic *BRCA* variants only among the sisters of TMM CommCohort members. The carriers found in the TMM CommCohort were mainly male and the female carriers were relatively young, so they themselves had not yet accumulated many cancer cases. A preponderance of a history of cancer in the mothers was not observed, but the mothers should have been aged more than 80, so their accumulation of sporadic cancers would have masked the HBOC cases. Our study suggests that the self-reported data of the TMM CommCohort are useful to analyze the genotype–phenotype relationships, at least in cancer-predisposing syndromes.

Many of the cancer-predisposing genes are known to be associated with juvenile cancer syndromes such as Li Fraumeni syndrome. The variants responsible for juvenile cancer syndromes are usually very pathogenic and show strong effects on gene functions. It is not so easy to estimate the clinical significance of the moderately pathogenic variants in those genes that may have clinically significant effects on the hosts’ predisposition for cancer, but these moderately pathogenic (or hypomorphic) variants are now classified as VUSs. The insight that VUSs may have moderate but significant effects on cancer onset that can be reduced by personalized health care based on the genetic variant information. Around 10 years ago, a review paper by Berger et al. proposed that haploinsufficiency is not so uncommon in the onset of cancer in HBOC patients with pathogenic variants in the *BRCA* genes [51]. In terms of precision medicine and/or personalized health care, moderately disease-contributing variants are also critical [52]. Such variants may not be critical to prompt for taking radical actions such as prophylactic surgery, but the carriers may be encouraged to continue close health checks to detect HBOC cancers as early as possible.

## Conclusions

The present study indicates that a large dataset of Japanese whole-genome sequencing data (3.5KJPNv2) includes definitely and potentially pathogenic variations in representative genes responsible for HBOC: *BRCA1* and *BRCA2*. ClinVar and the ACMG-guided annotation tool InterVar detected more than 20 variants as pathogenic or likely pathogenic, including one obviously benign variant. In addition, the use of the combination of computational scoring and MAF picked up another eight candidates, including one likely benign mutant as defined by ClinVar. Some of the variants show concordance with other databases in terms of the pathogenic annotations. The self-reported individual and family histories of the carriers of potentially pathogenic *BRCA* variants were analyzed and the carriers’ sisters showed a significant history of cancer themselves. This study indicates that prospective genomic cohort studies is a powerful tool for identification of pathogenic variants. The present study will be useful to identify such moderately disease-contributing variations in populations and will contribute to the development of personalized health care based on the individual genomic information

## Acknowledgment

We thank all past and present members of Tohoku Medical Megabank Organization at Tohoku University (present members are listed at https://www.megabank.tohoku.ac.jp/english/a191201/). We also thank Edanz (https://www.edanzediting.co.jp) for editing the English text of a draft of this manuscript.

## Ethics approval and consent to participate

The study has been approved by the ethics committee of Tohoku Medical Megabank Organization at Tohoku University (registration number: 2018-4-003). All the participants of the present study are recruited by Tohoku Medical Megabank Organization at Tohoku University with written informed consent for participation of the cohort study. The study project is approved by the Tohoku Medical Megabank Organization.

## Availability of data and material

The variants data can be downloaded from jMorp database (see methods). To protect the participants’ privacy, the self-report data of the TMM CommCohort participants is available after approval by the steering committee for Materials and Information of TMM project.

## Authors’ contributions

The grand design of this study was done by HT, YW, NY, and JY. Planning of data analysis was done by AH, SO, KI, and HT. Statistical analysis was done by KI and JY, with the help by AH, HT, and YYK. Cohort date acquisition and curation was done by YH, NF, AH, KK, and SO. Discussions for gynecological implications are done by SS, MS, HT, and NY. Genomic data has been obtained by ST, SI, FK, and KK. The manuscript was written by HT and JY. General coordination of this whole project was done by MY.

## List of Supporting Information

Table S1 InterVar P or LP variants in the BRCA genes of the RIKEN 2,234 Japanese whole genome sequence dataset

Table S2 Summary of candidate “pathogenic” variants of BRCA genes in 3.5KJPN version 2

Table S3 Details of “pathogenic” BRCA variants but not P or LP by InterVar in Momozawa et al. (Ref. 13)

Table S4 Statistics of the offspring cancer histories of candidate BRCA pathogenic variants carriers in the TMM CommCohort.

